# A Quantum Lens on Molecular Design: A Machine-Learned Energy Function from Interacting Quantum Atoms

**DOI:** 10.64898/2026.03.03.709242

**Authors:** M. Hoffmann, A. Kazimir, T. Oestereich, L. Kaermer, F. Engelberger, J. Meiler, C. Lamers

## Abstract

Accurate predictions of the interactions (covalent bonds and non-covalent contacts between atoms) in a molecular system **require scalable, accurate, and interpretable energy functions**. While classical force fields and knowledge-based energy functions struggle to capture key electronic effects, quantum chemistry approaches such as density functional theory (DFT) provide the necessary accuracy but remain computationally demanding. Furthermore, gaining insight into interactions requires energy decomposition schemes. The **Interacting Quantum Atoms (IQA)** scheme is exceptionally attractive, offering a chemically intuitive, electron density (ED) topologically based separation into intra- and interatomic contributions, however its **high computational cost remains a significant barrier** for application to larger systems or tasks like ligand screening in drug discovery.

We address these limitations by introducing a **novel machine learning (ML) framework to predict accurate energies derived from the IQA scheme together with a comprehensive dataset of molecular systems and their calculated IQA decomposed energies**. It enables the rapid and accurate prediction of DFT single point energies and dissects these energies in a physically meaningful and chemically intuitive manner. Our method predicts all **intra-atomic energies and inter-atomic interaction energies (covalent and non-covalent)** within a defined distance cutoff, providing an energy function that decomposes the total energy into specific atomic contributions. This advance makes the IQA method viable for analyzing interaction energies in applications previously inaccessible due to computational expense, such as elucidating ligand-binding mechanisms and informing rational drug design.

## Introduction

Accurate energy functions are essential for evaluating molecular stability, analyzing interfaces, and optimizing molecular geometries (Alford et al. 2021). As the energy function is highly dependent on representation of the variety interactions (covalent bonds and non-covalent contacts) in the molecular system, it is essential to find a way for their accurate description. While classical force fields have proven valuable in modelling the variety of interactions, they typically approximate the continuous charge distribution with localized discrete atomic point charges (Paton and Goodman 2009), (Monticelli and Tieleman 2013), (Singh et al. 2025). This simplification imposes a fundamental limitation, as it struggles to capture some crucial interactions dominated by electron delocalization, such as cation–π, CH–π, or halogen bond interactions, leading to a physically incomplete description of the molecular structure and its stability (Vitalini et al. 2015; Unke et al. 2021). Moreover, due to the absence of the interactions the empirical and statistical fitting included in many of their parameterizations to ‘‘compensate’’ the missing physics can introduce biases, such as favoring certain interaction types over others or spuriously stabilizing non-native conformations, which can lead to inaccurate design predictions (Kellogg et al. 2011; Rubenstein et al. 2018).

Achieving a truly predictive and physically meaningful energy function necessitates a direct accounting of the electron density (ED) distribution. Moving towards higher accuracy calculations, the field is increasingly turning to (quantum chemistry) QC methods, which are able to explicitly include electronic effects. While rigorous, wavefunction-based QC methods rely on a high-dimensional mathematical construct that is not physically observable, it is easier to operate with the methods based on three-dimensional ED. ED is a quantity of the spatial electron distribution that can be experimentally measured (using X-ray diffraction, neutron diffraction, electron scattering, etc.) (Tsirelson and Ozerov 2020) or obtained by theoretical QC methods, such as density functional theory (DFT) (S. Phipps et al. 2015; Goerigk et al. 2011). Furthermore, those methods can provide essential details on the interactions types between atoms.

In order to deepen the understanding of the interaction nature in molecular systems the diversity of energy decomposition schemes were developed. However, most of these methods rely on orbital localization and require optimized wave functions, typically obtained for equilibrium geometries (Su et al. 2014; Gimferrer et al. 2023; Phipps et al. 2015). Another major limitation for energy decomposition analysis is their computational demand and difficulty of interpretation. Therefore, bonding analyses in larger molecular systems so far typically rely on classical or (semi-)empirical approximations, making accurate quantification of covalent and non-covalent energy contributions challenging (Mansoor et al. 2025; Thapa and Raghavachari 2019).

Among the vast variety of decomposition schemes, the Interacting Quantum Atoms (IQA) method is particularly appealing due to its simple and conceptually straightforward real-space formulation. IQA decomposes the total molecular energy into intra-(monoatomic) and interatomic (diatomic) contributions based on Bader’s Quantum Theory of Atoms in Molecules (QTAIM), providing a chemically intuitive picture of atomic and bonding interactions without relying on orbital partitioning (Bader and Nguyen-Dang 1981). In IQA, the ED gradient lines identify the shape of an atom in the molecule (**Figure 1**). The borders of the atom (atom basin) are separated by the condition ∇ρ(**r**)n(**r**)=0 (zero flux, forming zero flux surface), which corresponds the space of the molecules, where the ED gradient lines of atom A connect with the ED gradient lines of atom B.

**Figure 1.**
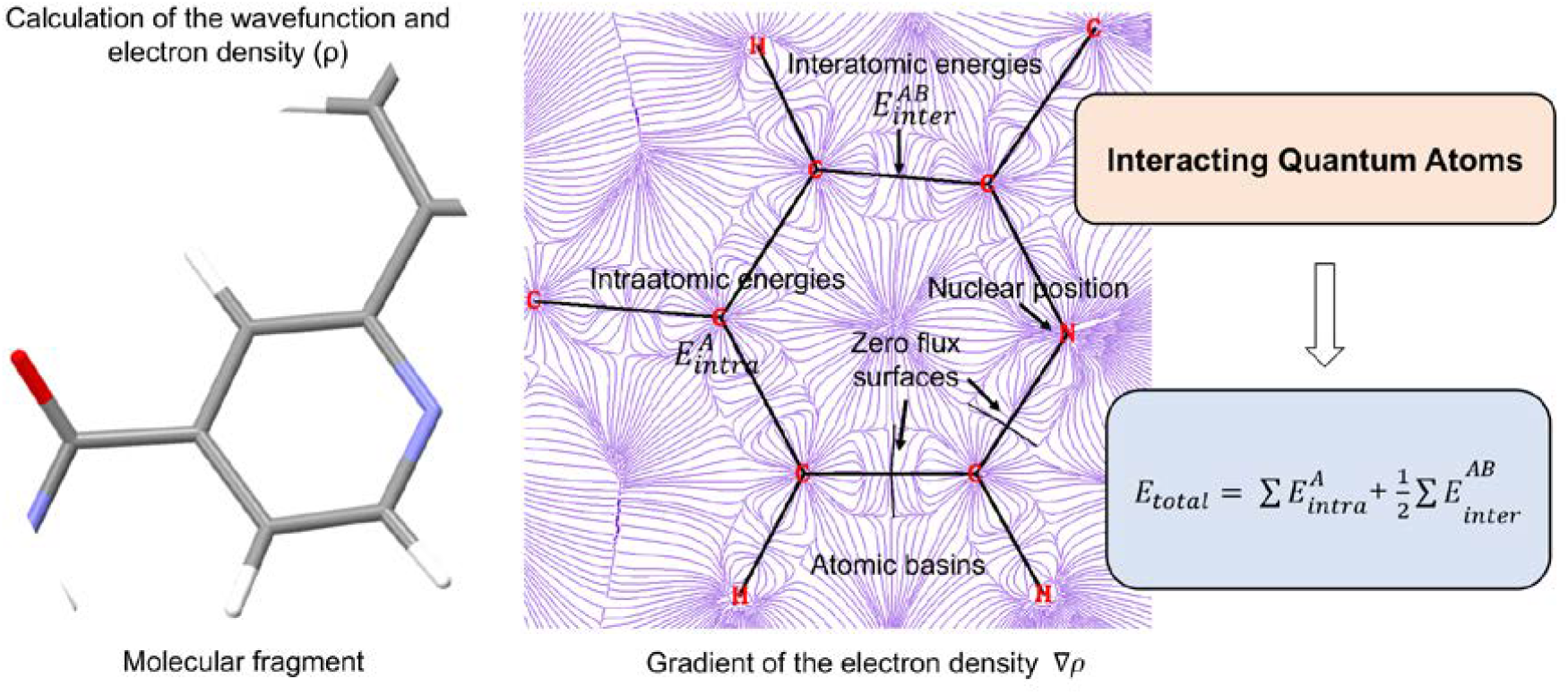
Interacting Quantum Atoms: The space of the molecule is decomposed using gradients of electron density.

The interaction appearing between two basins of atoms can be further characterized in terms of electrostatic, exchange and correlation contributions providing further details for chemical bonding (Guevara-Vela et al. 2020). Interestingly, IQA has been shown to recover energy components analogous to those obtained from traditional energy decomposition analysis schemes, while it additionally provides insights in the impact of the covalent bonds (Martín Pendás et al. 2006). As the IQA scheme uses heavy integration steps, it is hard to be applied for larger systems like proteins. By combining IQA with QM/MM (López et al. 2022), or accelerating IQA with the relative energy gradient (Falcioni and Popelier 2023), approaches have been proposed to tackle this challenge.

To bypass the high computational costs of QC calculations and make them accessible for larger scale applications in e.g. drug or protein discovery, machine learning (ML) approaches have been combined with the variety of QC methods (Schütt et al. 2018; Unke et al. 2021). Recent efforts have led to extensive datasets containing millions of molecular structures and their QM features like DFT single point energies and forces, as well to new types of novel architectures for deep neural networks (Smith et al. 2020). A significant development is the introduction of equivariant architectures capable of utilizing higher-order representations (Geiger and Smidt 2022). It enabled the network to encode complex geometric features analogous to atomic orbitals, addressing the limitations inherent in simple scalar or vector descriptions. A notable example is the Open Molecule 2025 Dataset (OM25 containing diverse fragments with up to 350 atoms including electrolytes and biomolecules), which has been used to train specifically designed equivariant neural networks on over 100 million structures. leading to state-of-the-art machine learning interaction potentials and energy functions (Levine et al. 2025).

Within the specific domain of IQA, pioneering works have already demonstrated the potential of ML to predict energy components and real-space chemical descriptors. For instance, the DL_FFLUX force field utilizes Gaussian process regression to learn atomic properties and IQA energies with quantum mechanical accuracy (Symons et al. 2021), while the SchNet4AIM architecture employs message-passing neural networks to accurately predict local atomic and interatomic IQA descriptors (Gallegos et al. 2024). However, these previous approaches rely on Gaussian process regression or invariant spatial descriptors and are typically trained on smaller datasets.

Therefore, to overcome the scalability barrier and achieve broader generalization we have developed a framework based on the **EquiformerV2 equivariant graph neural network** that learns the IQA energy decomposition across a dataset covering more than 130’000 diverse molecular structures. Our **ML-IQA** framework provides quantum-mechanically accurate **intra- and inter-atomic energy** components at a fraction of the traditional computational cost, thereby transforming IQA into a tool for analyzing interaction energies in applications previously inaccessible due to computational expense. This approach further will also allow decomposition of these energies in a **physically meaningful and interpretable** manner. Specifically, our model yields intra-atomic energies and interatomic **covalent and non-covalent interaction** energies for every atom and atom pair within a defined spatial range. The resulting detailed energy breakdown transforms our model into a tool for molecular analysis, offering insights into the stabilizing and destabilizing energy contributions within complex molecular systems, making it a building block for analyzing interaction patterns, approximating binding enthalpies and informing rational drug design within a broader thermodynamic framework (Combs et al. 2013).

### Experimental Section

#### Interacting Quantum Atoms Scheme

The Interacting Quantum Atoms (IQA) approach is founded on the Quantum Theory of Atoms in Molecules (QTAIM) (Guevara-Vela et al. 2020). In this framework, the gradient field of the electron density (ED) partitions molecular space into distinct atomic basins.

These basins are bounded by interatomic surfaces on which the gradient vectors of the ED are everywhere perpendicular to the surface and have zero flux across it. In IQA scheme the entire energy of the molecule can be decomposed into monoatomic intra- and diatomic interatomic contributions (eq. 1):

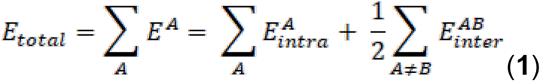

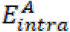 as it is the intraatomic contribution (‘‘self-atomic’’ energy component) of an atom (atom A),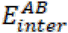 while refers to the diatomic interaction term between a couple of atoms (atom A and atom B). The energy terms calculated for the basin of atom A can be further decomposed into intra- and interatomic contributions according to equations (2) and (3):

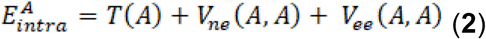

These are the terms which are identified as kinetic energy of electrons in basin of atom A (*T(A)*), attraction of electrons of atom A to the nucleus of atom A (*Vne(A,A)*) and electron-electron interaction in basin A (*Vee(A,A)*). While interatomic term can be expressed as:

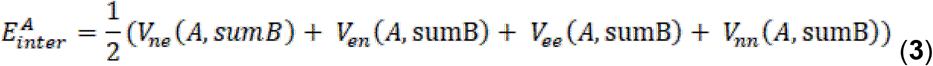

In equation (**3**) the interaction terms represent the coupling of atom A with the nuclei and electron densities of all other atoms in the molecular system, excluding self-interactions within A (*Vne(A, sumB), Ven(A, sumB), Vee (A, sumB)*). In addition, the nuclear–nuclear repulsion between the nucleus of atom A and the nuclei of the other atoms in the molecule is also included (*Vnn(A, sumB)*).

The diatomic terms will contain the next components in equation 4, where B is denoted as any other atom in the molecular system:

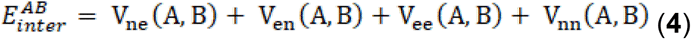

Additionally, in equations (2)–(4), the electron–electron interaction terms can be further decomposed into exchange (*VeeX*) and correlation (*VeeC*) components, which represent valuable quantities both for physical interpretation and as parameters to be used for machine learning.

### Note

In AIMAll output, the interaction of a given atom A with the rest of the molecule can be reported in two formally equivalent ways: as the sum of all pairwise interatomic interactions, *E*_inter_ (*A,sumB*), or as the interaction of A with the remainder of the molecule,*E*_inter_ (*A, A*′) = *E*_inter_(*A, Mol*)−*E*_inter_(*A, A*), where A′ denotes the union of all other atomic basins. Both definitions are theoretically identical,so *E*_inter_ (*A,sumB*) ≈ *E*_inter_(*A, A*′); however, small numerical differences may appear in practice due to finite integration grids and rounding errors. AIMAll always reports the quantities *E*_inter_(*A,sumB*) and *E*_inter_(*A, A*′) and the difference between them, as a diagnostic measure of this integration accuracy, the latter ideally should be close to zero.

### Dataset Generation and Curation

The training dataset was constructed by combining structures from several established quantum chemistry datasets to ensure broad coverage of chemical space and interaction types. These include A24 (Řezáč and Hobza 2013), Ionic H-Bonds (Řezáč and Hobza 2012), JSCH-2005 (Paton and Goodman 2009), NCI Atlas (Řezáč 2020b, 2020a; Kříž et al. 2021; Kříž and Řezáč 2022; Řezáč 2022), OMol25 (Levine et al. 2025), S26-extra (Riley and Hobza 2007), S66x8 (Řezáč et al. 2011), SCAI (Berka et al. 2009), Small Halogen Bonding Complexes (Riley et al. 2009), Water Clusters (Bryantsev et al. 2009) and X40x10 (Řezáč et al. 2012). The atom types were limited to H, C, N, O, S, P, Cl, and F to find a balance between computational cost, complexity of the learning task and covering the important atom types present in biomolecules. For model validation and testing, we utilized a dataset of isolated small peptides containing aromatic side chains (Valdes et al. 2008), which were held out from the training data. The final combined dataset comprises a total of 131,714 molecular structures, including charged structures and a high diversity of non-equilibrium structures to encourage the model to learn a highly accurate potential energy surface and avoid collapse outside of equilibrium points (**Figure 2**) (Smith et al. 2017).

**Figure 2.**
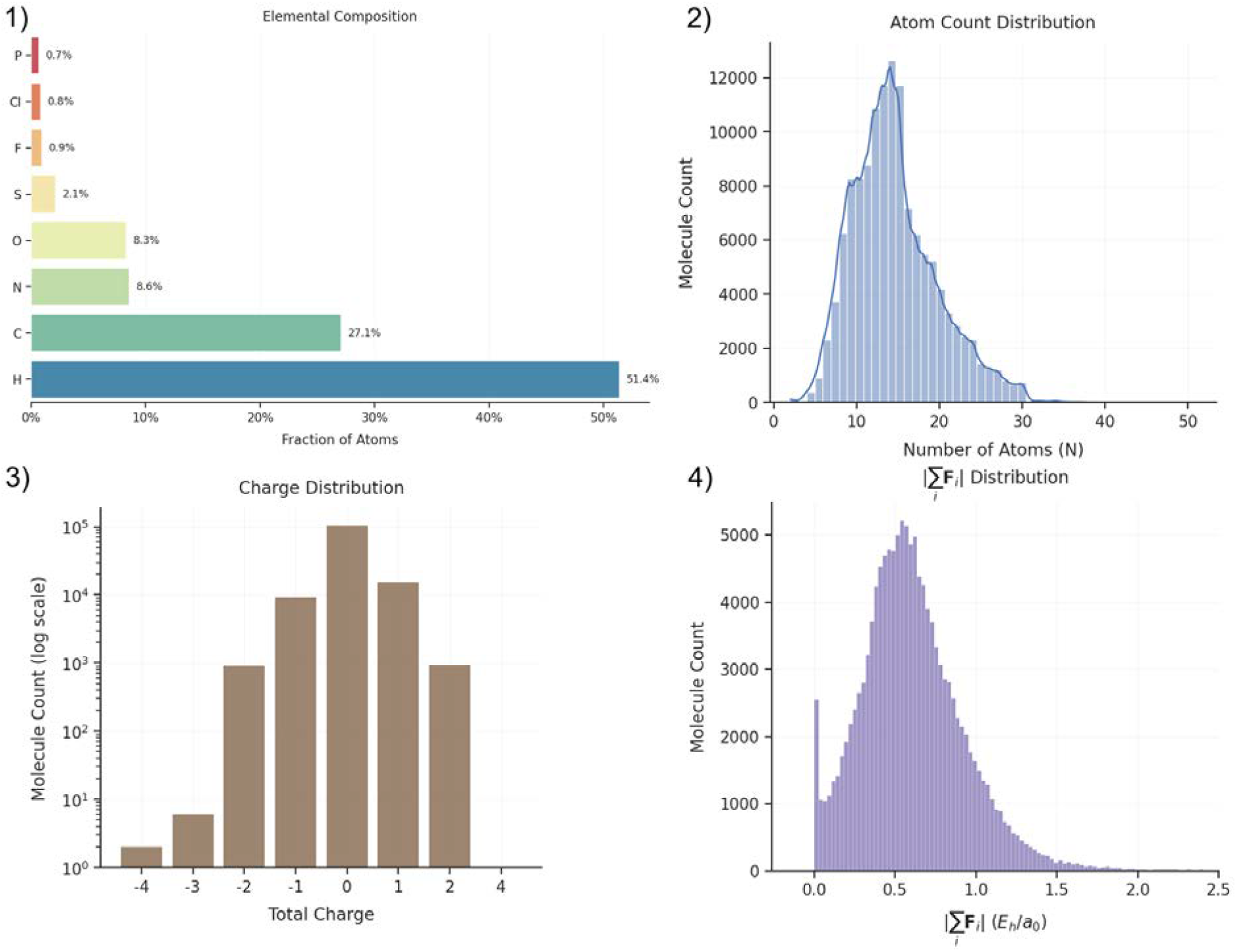
Dataset statistics. 1) displays the percentage-wise contribution of each atom type to the dataset, 2) the distribution of molecular sizes and 3) the variety of differently charged molecules included. 4) shows the distribution of the resultant force norm *(Hartrees per Bohr)* across the atoms *i* for each molecule. Structures close to the equilibrium have a sum of forces acting on their atoms close to zero.

For each structure, the electronic wavefunction was calculated using the Gaussian 16 software package (Frisch et al. 2016). The following IQA decomposition was performed using the AIMAll software (Keith 2019). The wave function was obtained at M06-2X/ jun-cc-pVDZ level of theory (Zhao and Truhlar 2008). To capture NCIs properly we have chosen M06-2X which includes the dispersion correction, also important for long-distance and weak interactions. **Importantly**, AIMAll supports few functionals including M06-2X, where the exchange-correlation energy term (VeeX) is a part of the functional, otherwise it approximates this term as Hartree-Fock exchange functional. Its empirical fitting of the functional parameters provides the most substantial correction for dispersion effects, however no DFT functional with the additive dispersion corrections can be included (Hohenstein et al. 2008; Goerigk et al. 2011). The jun-cc-pVDZ basis set of the Dunning hierarchy was chosen for our calculations to provide a favorable balance between computational cost and the accurate description of non-covalent interactions, such as dispersion effects. As a “Calendar” basis set, it comprises polarizable and diffuse functions and allows, simultaneously, the removal of high-angular momentum diffuse functions to mitigate over-complexity in our computational setup (Papajak et al. 2011).

To ensure the integrity of the ground-truth data, a filtering step was implemented after the calculations were complete. We established a maximum tolerance for the mismatch between the single point total energy of the molecular complexes obtained by the DFT calculations and the total energy obtained from the sum of the atom-wise energy obtained by IQA decomposition, set to 0.001 Hartrees (Eh) per molecule (**Figure 3**). Any structures exceeding this threshold were removed from the dataset to filter out inaccurate data points that may have arisen from integration errors within the IQA procedure (**Figure 4**).

**Figure 3.**
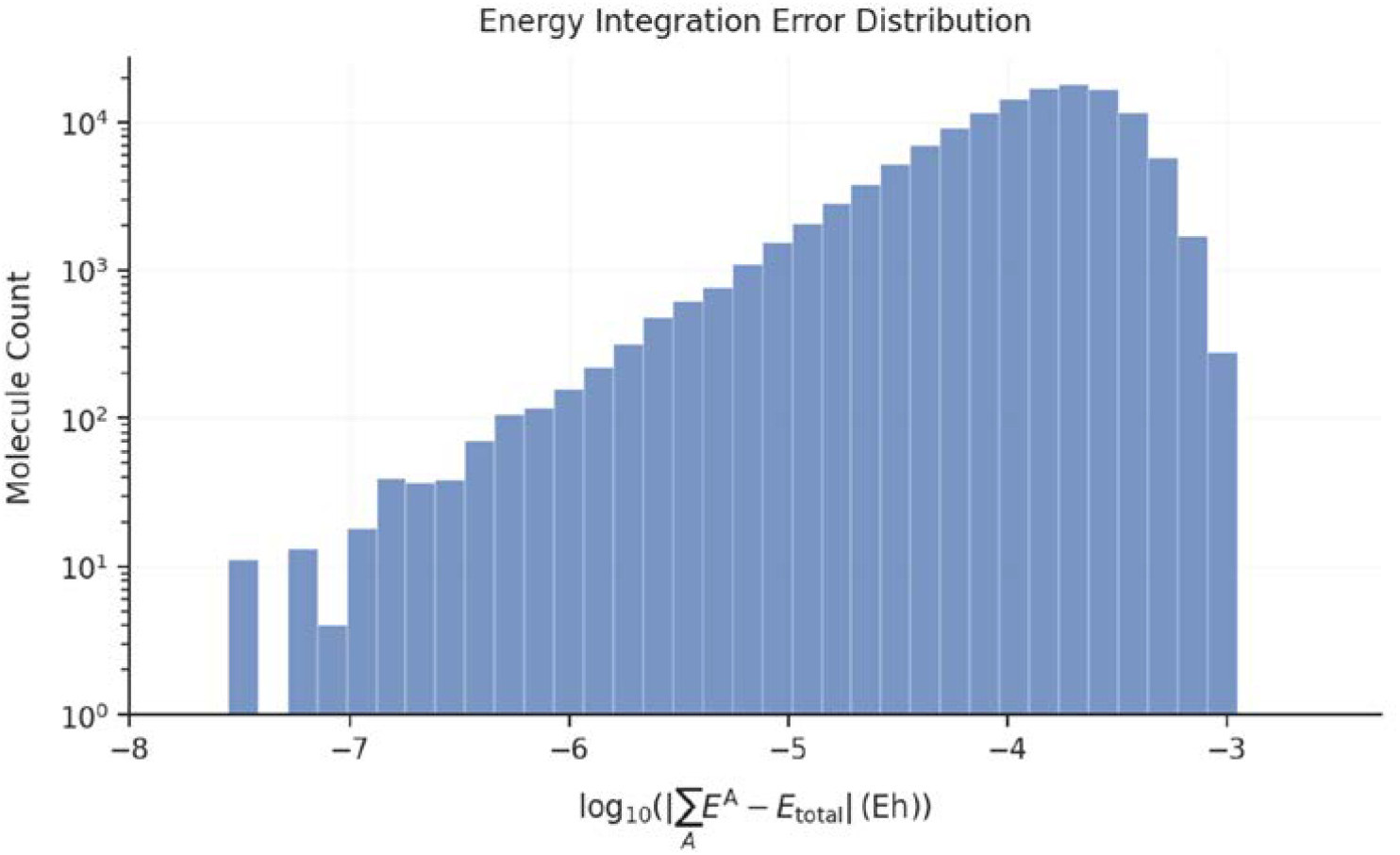
Integration error across the dataset. The distribution of the remaining integration error in the final dataset, defined as the mismatch between the sum over the atomic energies for a molecule and its single point DFT energy.

**Figure 4.**
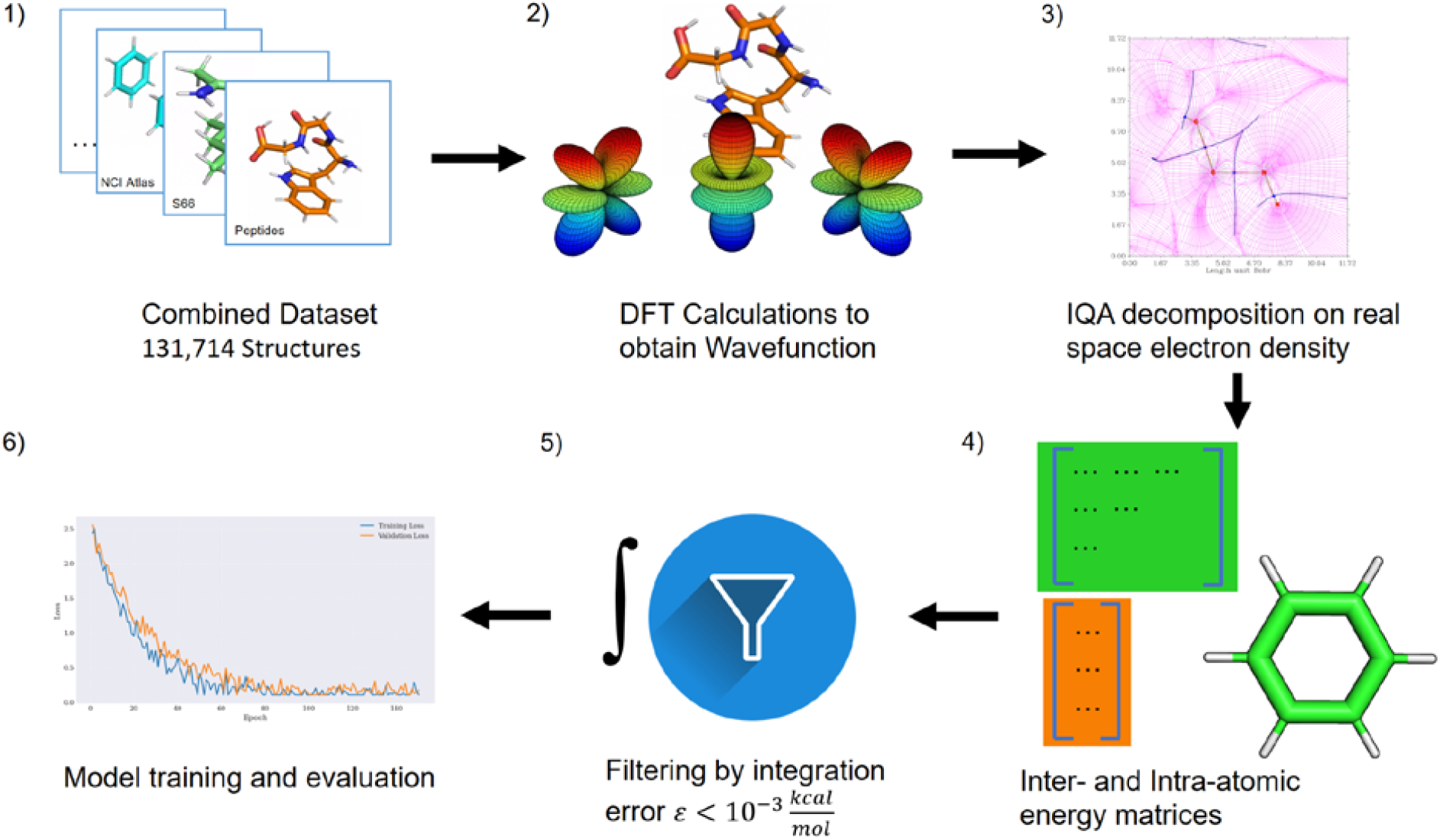
Workflow scheme for data generation and processing. 1) Collecting molecular structures from different dataset sources to create a rich and diverse dataset for the energy prediction task. 2) DFT calculations (M062X/jun-cc-pVDZ) were performed on the geometries in the obtained dataset, producing the wavefunctions for these molecular systems. 3) Integration over atomic basins of the real space ED for obtaining the decomposed IQA energies. 4) IQA energy dataset creation; packing the intra- and inter-atomic IQA energy matrices for data export and further machine learning model production. 5) To obtain the final version of the IQA dataset, the pre-version was filtered by the integration error of the calculations for each molecular system. Due to the numerical integrations in step 3, the sum of the decomposed energy for a molecule varies from the value of the DFT energy of the wavefunction in step 2. Molecules where the absolute value of the difference between DFT energy and sum of the IQA terms was greater than 0.001Eh (0.63 kcal/mol) were removed from the dataset. 6) The IQA model was trained on the final dataset.

### Model Architecture

The predictive model is built upon an equivariant graph neural network, specifically using the EquiformerV2 architecture as its core component. It is built upon the concept of the E3NN package (Geiger and Smidt 2022) and extends it functionality by faster and more efficient equivariant feature mapping and processing as it was also the top performing model and state of the art architecture (state 2024) for various tasks of the Open Catalyst Challenge (Liao et al. 2024; Sriram et al. 2024). Inherent SE(3)-equivariance allows the model to respect rotational and translational symmetries of molecular systems, improving physical consistency and generalization, a widely adopted and effective property in modern physics-informed ML potentials. The EquiformerV2 allows for higher-order equivariant representations defined by the maximum degree of spherical harmonics, *l*_*max*_. This parameter controls the angular resolution of the atomic features, effectively allowing the network to model complex, anisotropic geometric information that simple scalar or vector representations cannot capture, akin to atomic orbitals (e.g., d or f orbitals).

This base architecture was enhanced with a custom prediction head designed to output the decomposed IQA energy terms. It enables the prediction of node- and edge-wise scalar values for the energy prediction regression task. Within this framework, intra-atomic features are predicted as node properties, and inter-atomic features are treated as edge properties. The model operates on a radius graph defined by a distance cutoff, aggregating local information via message passing. Direct interatomic energies are predicted exclusively for edges within this graph. However, the stacking of multiple layers expands the reception of a node beyond the cutoff, capturing long-range dependencies and allowing for predicting the sum of the all interactions for an atom as a node feature. The loss function for the model is defined as the Mean Absolute Error (MAE) between the ground-truth and predicted values for every individual term of the IQA decomposition scheme (**Figure 5**).

**Figure 5.**
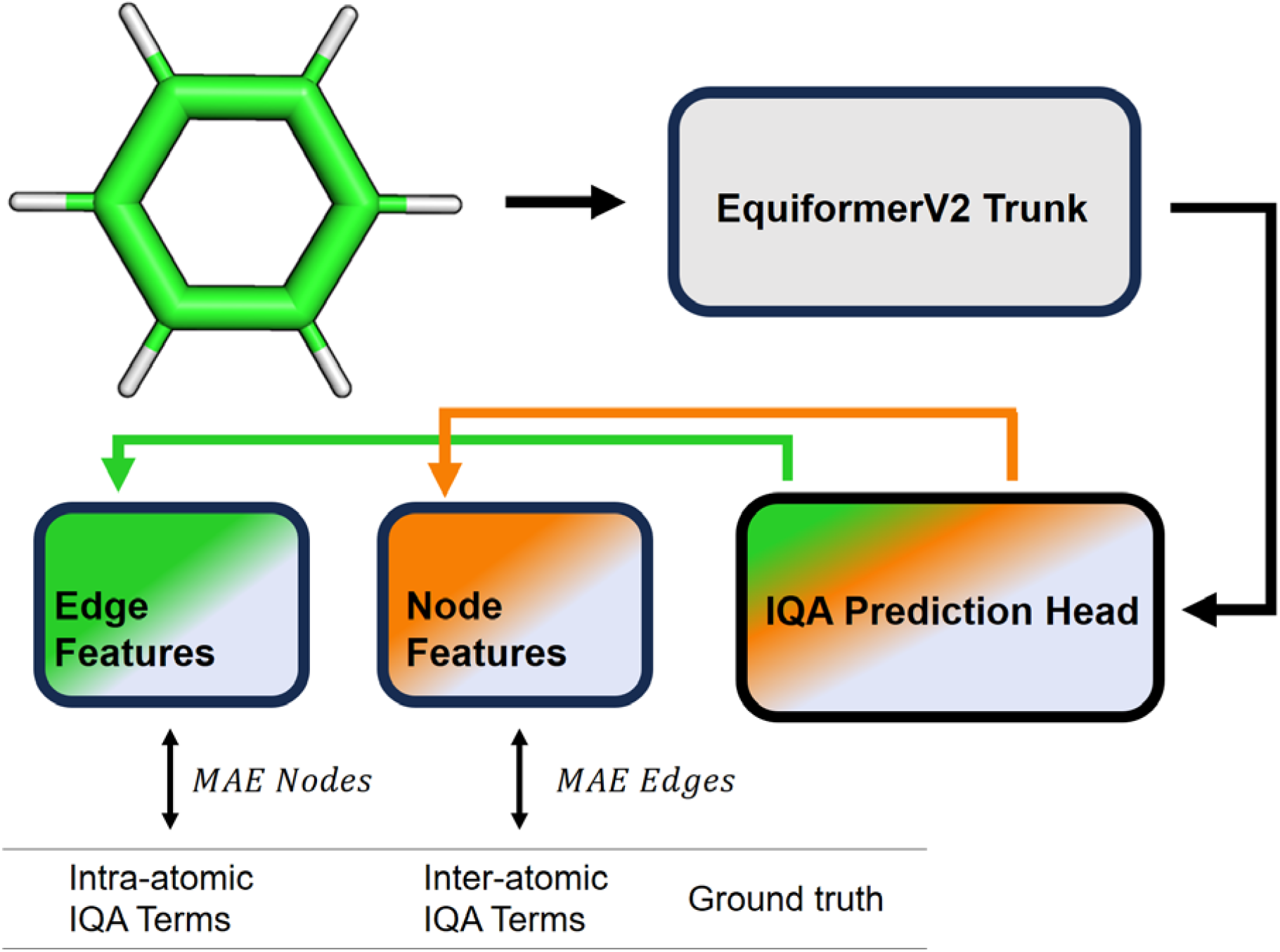
Model architecture overview. The molecular geometries consisting of atomic coordinates and atom types serve as input features for the IQA model. The EquiformerV2 architecture serves as the model’s trunk, calculating the node and edge embeddings. The IQA Prediction Head is integrated atop the Equiformer core, designed to predict the node- and edge-wise IQA energies. The loss, defined as the mean absolute error (MAE) is then calculated by comparing the node and edge predictions with the monoatomic and diatomic energies from the dataset respectively.

In this work, we focused on building the first prototype of an ML-IQA model and trained specifically on the total atom-wise energy and the atom pair-wise energy The key hyperparameters defining the model architecture are provided in Table 1.

**Table 1:** Model Hyperparameters. *max_distance* defines the cut-off value for the radius graph of the model, in atomic units. Beyond this cutoff, interactions between two distinct atoms are not predicted directly. *num_layers* defines the number of stacked message passing layers in the model’s trunk. *l_max* defines the maximum degree of spherical harmonics.

| Parameters | Value |
| --- | --- |
| **max_distance** | **12.0 a.u.** |
| **num_layers** | **6** |
| **l_max** | **5** |

#### Training Protocol

The models were trained for 100 epochs, until the validation loss stopped decreasing, using a batch size of 32. A CosineAnnealing learning rate schedule was employed, with an initial learning rate of 1e-4 decaying to a final target rate of 1e-5. The specific parameters for the training procedure are detailed in Table 2.

**Table 2:** Training Hyperparameters. The model was trained in 100 epochs with a batch size of 32. A cosine annealing learning rate scheduler was employed using the defined initial and target learning rate.

| Parameter | Value |
| --- | --- |
| **epochs** | **100** |
| **batch size** | **32** |
| **learning rate schedule** | **CosineAnnealingLR** |
| **initial learning rate** | **$1e-4$** |
| **target learning rate** | **$1e-5$** |

## Results

To develop a ML-IQA framework, we trained on the prediction of the total energy term (*E*^*A*^, see equation (2)) and the diatomic interatomic energy (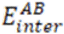, see equation (4)) per atom. The model was evaluated a dataset of small peptides with aromatic side-chains (Valdes et al. 2008) as well as on the S22x5 dataset (see **Appendix**).

The model achieves a mean absolute error (MAE) of 1.65×10^-4^ Eh (0.10 kcal/mol) for *E*_*inter*_*(A,B)/2* on the peptide test set. Overall, the model’s predictions of the interaction energies correlate well to the ground-truth (**Figure 6** 2)). When plotting the absolute errors against the reference interaction energies, we observe that for the majority of the interactions - which are non-covalent, weak interactions - the model’s predictions have a very low error. For covalent interactions, the magnitude of the energies themselves are significantly higher, explaining a relatively larger error margin in this region (**Figure 6** 4)). The model accurately captures distinct clustering patterns within the covalent bonding range of −0.7 Eh to −0.4 Eh (single, double, triple bonds), demonstrating its ability to differentiate between these various covalent bond types by also predicting their energies accurately (**Figure 6** 4)). For the *E*_*intra*_*(A)* energy the MAE of the model is evaluated at 8.87×10^-3^ Eh (5.57 kcal/mol). A large fraction of the atomic energies is found close to zero, representing the hydrogen atoms in the molecular structures of the dataset.

**Figure 6.**
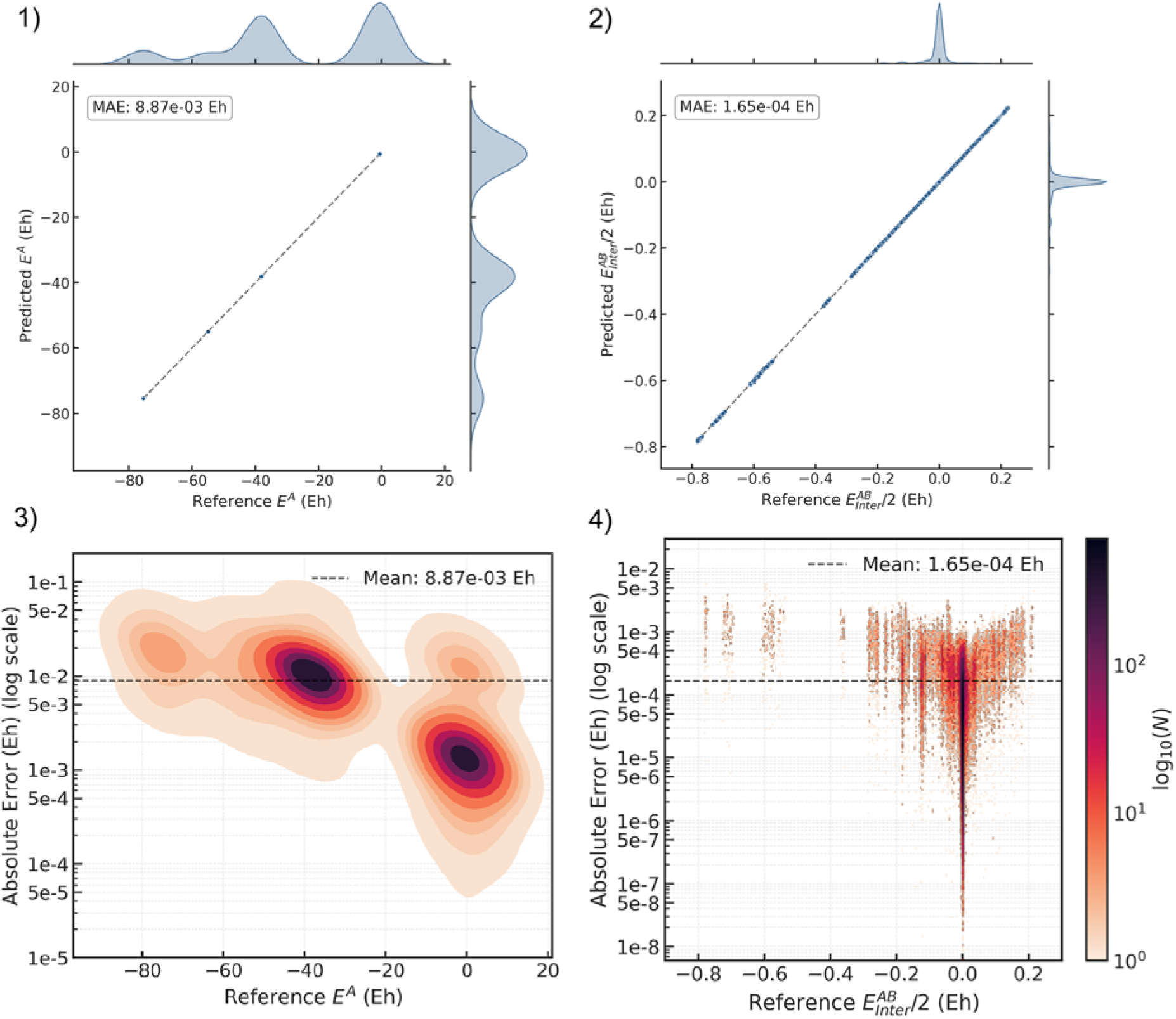
Evaluation of the IQA model on small peptides with aromatic side-chains. (Valdes et al. 2008). 1) Predicted per-atom total energy *E*_*intra*_*(A)* vs reference (ground truth) values. The predictions correlate with the true energy values with a mean absolute error (MAE) of 8.87×10^−3^ Hartrees (Eh) (5.57 kcal/mol). 2) Predicted atom pair-wise interaction energy *E*_*inter*_*(A,B)/2* versus reference (ground truth) values. The plot shows correlation between prediction and ground truth with a MAE of 1.65×10^−4^ Eh (0.10 kcal/mol). The interaction energies are predominantly near zero because high-energy covalent interactions represent a small fraction of all possible atomic interactions evaluated within the specified range. 3) MAE of *E*_*intra*_*(A)* in Eh and 4) MAE of *E*_*inter*_*(A,B)/2* in Eh plotted versus their corresponding ground truth energies. Darker colors represent agglomeration of atoms in this area of the plot.

To visualize the interactions between atomic pairs, atoms were connected by lines representing stabilizing (blue) or destabilizing (red) interactions. In **Figure 7**, two representative examples from the dataset of small peptides illustrate different types of non-covalent contacts for visualization purposes: (A) Tryptophan-Glycine (WG), highlighting hydrogen bonding and π-interactions involving the aromatic nitrogen, and (B) Glycine-Phenylalanine-Alanine (GFA), demonstrating π interactions involving the aromatic system and hetero-atoms outside the aromatic ring. For both the standard hydrogen bonds and the more complex aromatic interactions, the model accurately predicts the inter-atomic terms with deviations from the ground truth generally well below 0.001 Eh (0.63 kcal/mol). It should be noted that these predicted inter-atomic terms represent absolute, pairwise local interactions and are therefore not directly comparable to standard macroscopic hydrogen bond energies. The interactions predicted by the model are in good agreement with the ground-truth IQA results, demonstrating that the machine learning framework can reproduce the essential features of the computationally demanding IQA scheme at a fraction of its cost (**Table 3**).

**Table 3:** Computational cost comparison between numerical IQA calculations and the machine learning model (ML-IQA). The numerical IQA calculations include obtaining the wave function at M062X/jun-cc-pVDZ level of theory, integration of the wavefunction square to obtain the ED, defining topology of the ED and atomic basins, integration of the atomic basins to determine IQA energies.

| X-40 Dataset | Methane-Cl2 | HF-Methanol | Methanol-Chloromethane |
| --- | --- | --- | --- |
| Number of atoms | 7 | 8 | 11 |
| CPU *hours* numerical calculation | 4.40 | 0.90 | 18.35 |
| CPU *seconds* machine learning model | 0.019 | 0.029 | 0.049 |
| Speed up on CPU | $0.83 \times 10^6$ | $0.11 \times 10^6$ | $1.35 \times 10^6$ |
| **Small Peptides Dataset** | **GGF** | **GFA** | **WGG** |
| Number of atoms | 37 | 40 | 41 |
| CPU *hours* numerical calculation | 181.31 | 105.50 | 140.05 |
| CPU *seconds* machine learning model | 0.320 | 0.293 | 0.404 |
| Speed up on CPU | $2.04 \times 10^6$ | $1.31 \times 10^6$ | $1.26 \times 10^6$ |

**Figure 7.**
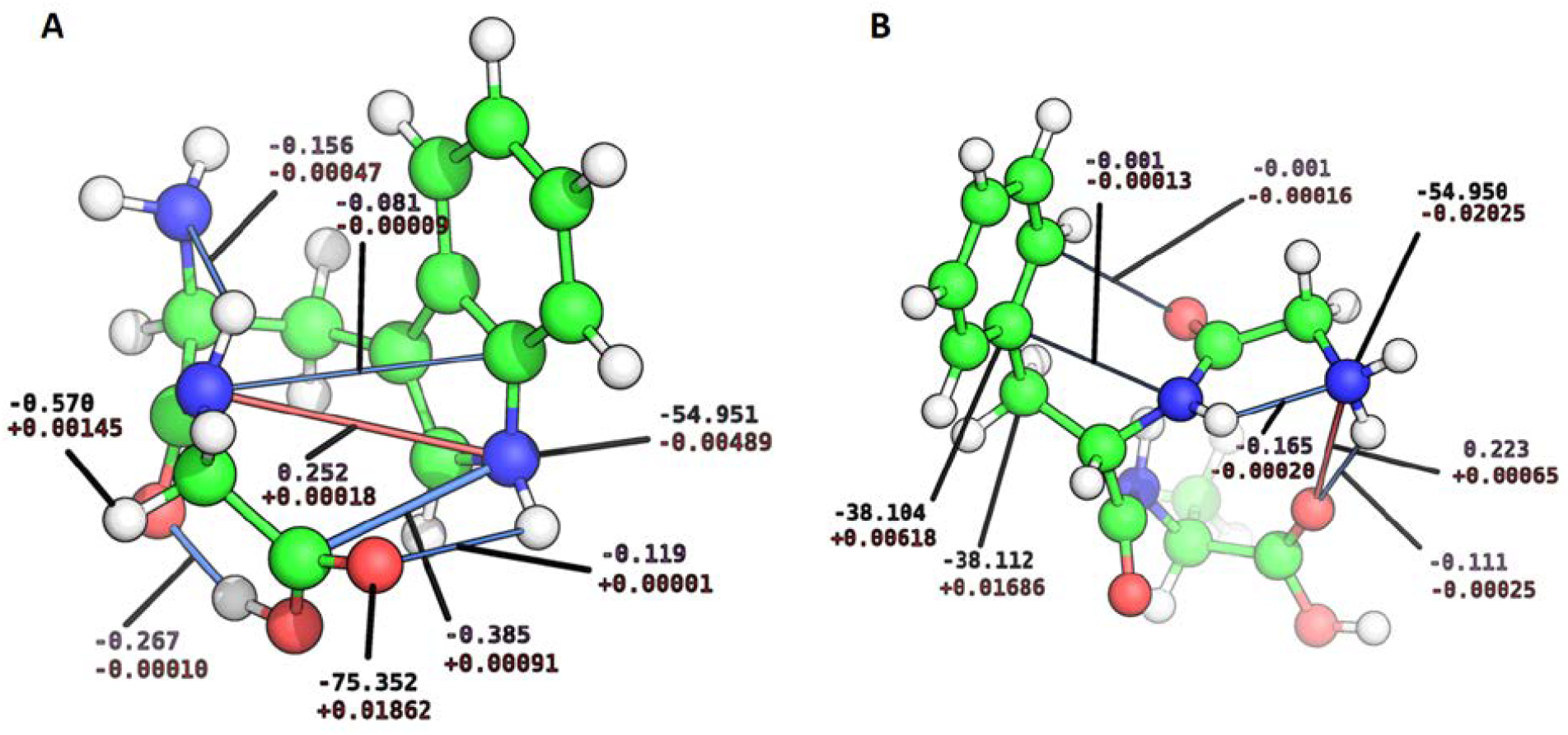
Exemplary non-covalent interactions for visualization purposes are shown as the lines connecting interacting atoms in the small peptides (A) Tryptophan-Glycine (WG) and (B) Glycine-Phenylalanine-Alanine (GFA), as predicted by the model. The complexes were chosen to represent different non-covalent contacts, including hydrogen bonding and π-interactions involving the aromatic nitrogen in (A), and π interactions involving the aromatic system and hetero-atoms outside the aromatic ring in (B). The stabilizing interactions (negative energy values) are shown as blue lines and destabilizing contacts (positive values) are shown as red lines. The strength of the interaction is additionally indicated by the thickness of each line. Predicted intra-atomic energies are labeled in black, while predicted inter-atomic interaction energies are labeled in violet. Below these predicted values, the difference compared to the ground truth is displayed in red. All values depicted have Hartree energy units.

We evaluated the computational performance of the ML-IQA model by comparing its execution time with that of the numerical IQA scheme. Both numerical IQA and ML-IQA calculations were performed on a CPU (AMD(R) EPYC(R) 7713 @ 2.0GHz). The resulting speed-up, reaching several orders of magnitude, highlights the computational efficiency of the ML-IQA model (see **Table 3**).

## Conclusions

We have demonstrated that an equivariant graph neural network can learn the IQA decomposition for the test dataset of small peptides, with MAEs of 8.87×10^-3^ Hartrees (Eh) (5.57 kcal/mol) for the total atom-wise energies and 1.65×10^-4^ Eh (0.10 kcal/mol) for the total diatomic interaction energies, showing its promise for scalable, interpretable molecular energy predictions. This work represents a step toward making the IQA method and its detailed chemical insights more accessible for tasks where direct calculation is computationally costly.

The low error for the interaction energies indicates the model is well-suited for the analysis of molecular interactions and for providing quantitative estimates of the atomic contributions that stabilize or destabilize a given conformation. However, the prediction error for the atomic-wise energies accumulates when summed, resulting in a total molecular energy prediction that is currently too inaccurate for use in applications that rely on precise energy gradients, such as molecular dynamics simulations (around 0.04 kcal/mol per atom for state-of-the-art machine learning interatomic potentials (Levine et al. 2025) in contrast to our 4.61 kcal/mol per atom MAE).

The IQA framework provides essential information for chemically intuitive interpretation and machine learning. It offers quantitative insight into molecular energy decomposition at the level of atoms. This approach allows for a detailed energy breakdown by atom or atom pair, which can be extended to the molecular fragments such as residue and residue-pair scores in protein systems.

Future work will focus on refining the model to increase the accuracy of the predictions by refining the model’s architecture, integrating data normalization techniques and increasing the training steps and to apply it on larger biomolecular systems and validating it for this use-case. Additionally, we will add a force prediction head for training and benchmark on atomic forces to improve the quality of the learned energy landscape of the IQA model and make it usable for molecular design and optimization tasks. Applying the IQA method to challenging tasks in drug design could improve the quality of *in-silico* models by analyzing for certain types and strength of interactions for instance in the case of ligand-target binding or interactions between substrate and catalytic site in enzyme design. We will also aim to incorporate a fully resolved decomposition framework including both electronic and nuclear terms, allowing deeper analysis of electronic interactions and their mechanisms.

