## Supplemental Figure 1 for "A Quantum Lens on Molecular Design: A Machine-Learned Energy Function from Interacting Quantum Atoms"

### Appendix

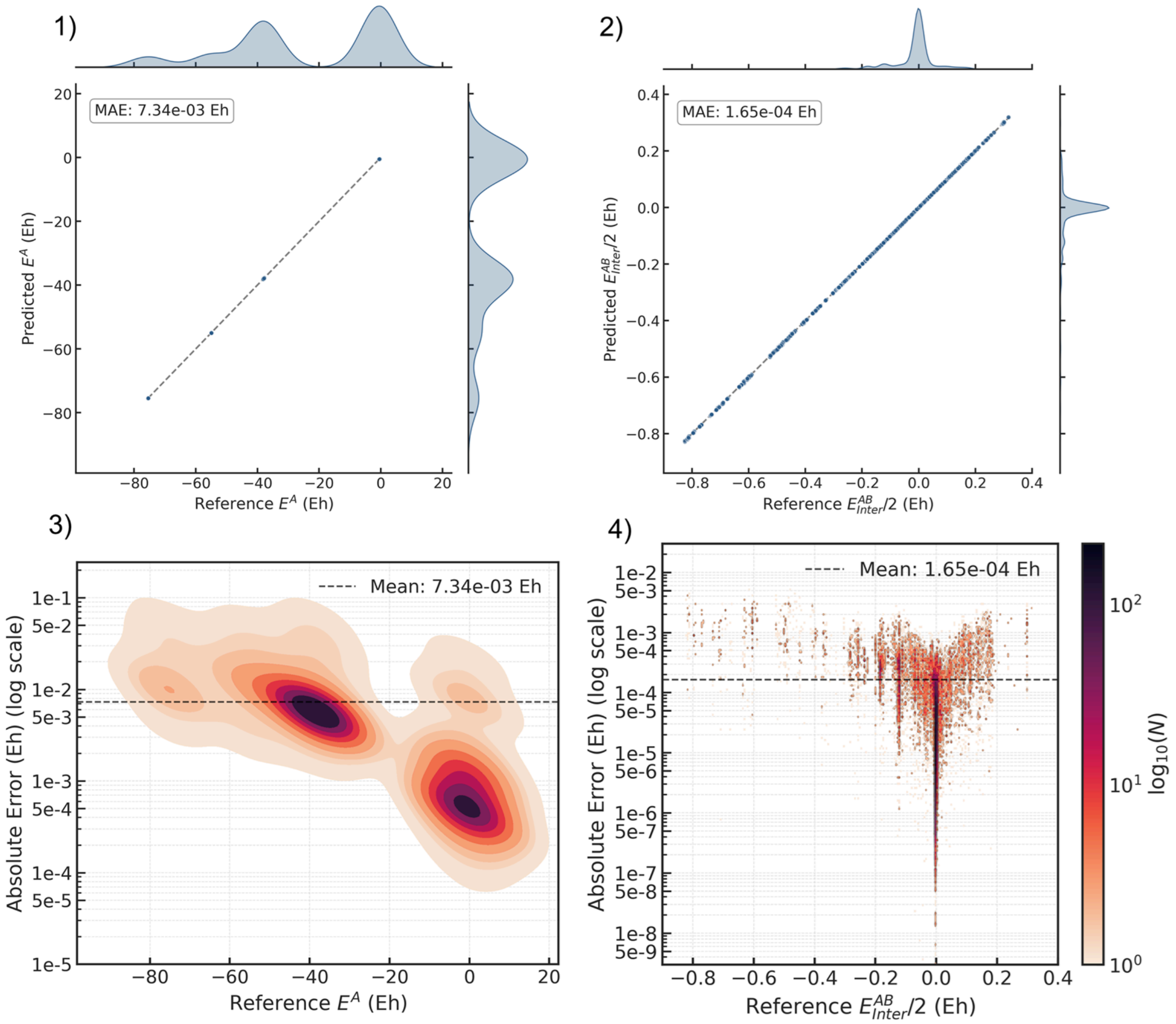


**Figure A1. Evaluation of the IQA model on the S22x5 dataset.** 1) Predicted per-atom total energy $E_{intra}(A)$ vs reference (ground truth) values. The predictions correlate with the true energy values with a mean absolute error (MAE) of $7.34\times10^{-3}$ Hartrees (Eh) (4.61 kcal/mol). 2) Predicted atom pair-wise interaction energy $E_{inter}(A,B)/2$ versus reference (ground truth) values. The plot shows correlation between prediction and ground truth with a MAE of $1.65\times10^{-4}$ Eh (0.10 kcal/mol). 3) MAE of $E_{intra}(A)$ in Eh and 4) MAE of $E_{inter}(A,B)/2$ in Eh plotted versus their corresponding ground truth energies. Darker colors represent agglomeration of atoms in this area of the plot.
